# Modular lipid nanoparticle platform technology for siRNA and lipophilic prodrug delivery

**DOI:** 10.1101/2020.01.16.907394

**Authors:** Roy van der Meel, Sam Chen, Josh Zaifman, Jayesh A. Kulkarni, Xu Ran S. Zhang, Ying K. Tam, Marcel B. Bally, Raymond M. Schiffelers, Marco A. Ciufolini, Pieter R. Cullis, Yuen Yi C. Tam

## Abstract

Successfully employing therapeutic nucleic acids, such as small interfering RNA (siRNA), requires chemical modifications or the use of nanocarrier technology to prevent their degradation in the circulation and to facilitate intracellular delivery. Lipid nanoparticles (LNP) are among the most advanced nanocarriers culminating in the first siRNA therapeutic’s clinical translation and approval. At the same time, their applicability as modular platform technology due to the interchangeable building blocks and siRNA payload hallmarks one of LNPs’ major advantages. In addition, drug derivatization approaches to synthesize lipophilic small molecule prodrugs enable stable incorporation in LNPs. This provides ample opportunities to develop combination therapies by co-encapsulating multiple therapeutic agents in a single formulation. Here, we describe how the modular LNP platform can be applied for combined gene silencing and chemotherapy to induce additive anti-cancer effects. We show that various lipophilic taxane prodrug derivatives and siRNA against the androgen receptor, a prostate cancer driver, can be efficiently and stably co-encapsulated in LNPs. In addition, we demonstrate that prodrug incorporation does not affect LNPs’ gene silencing ability and that the combination therapy induces additive therapeutic effects *in vitro*. Using a double-radiolabeling approach, we quantitively determined the LNPs’ and prodrugs’ pharmacokinetic properties and biodistribution following systemic administration in tumor-bearing mice. Our results indicate that co-encapsulation of siRNA and lipophilic prodrugs into LNPs is an attractive and straightforward approach for combination therapy development.

**GRAPHICAL ABSTRACT:** 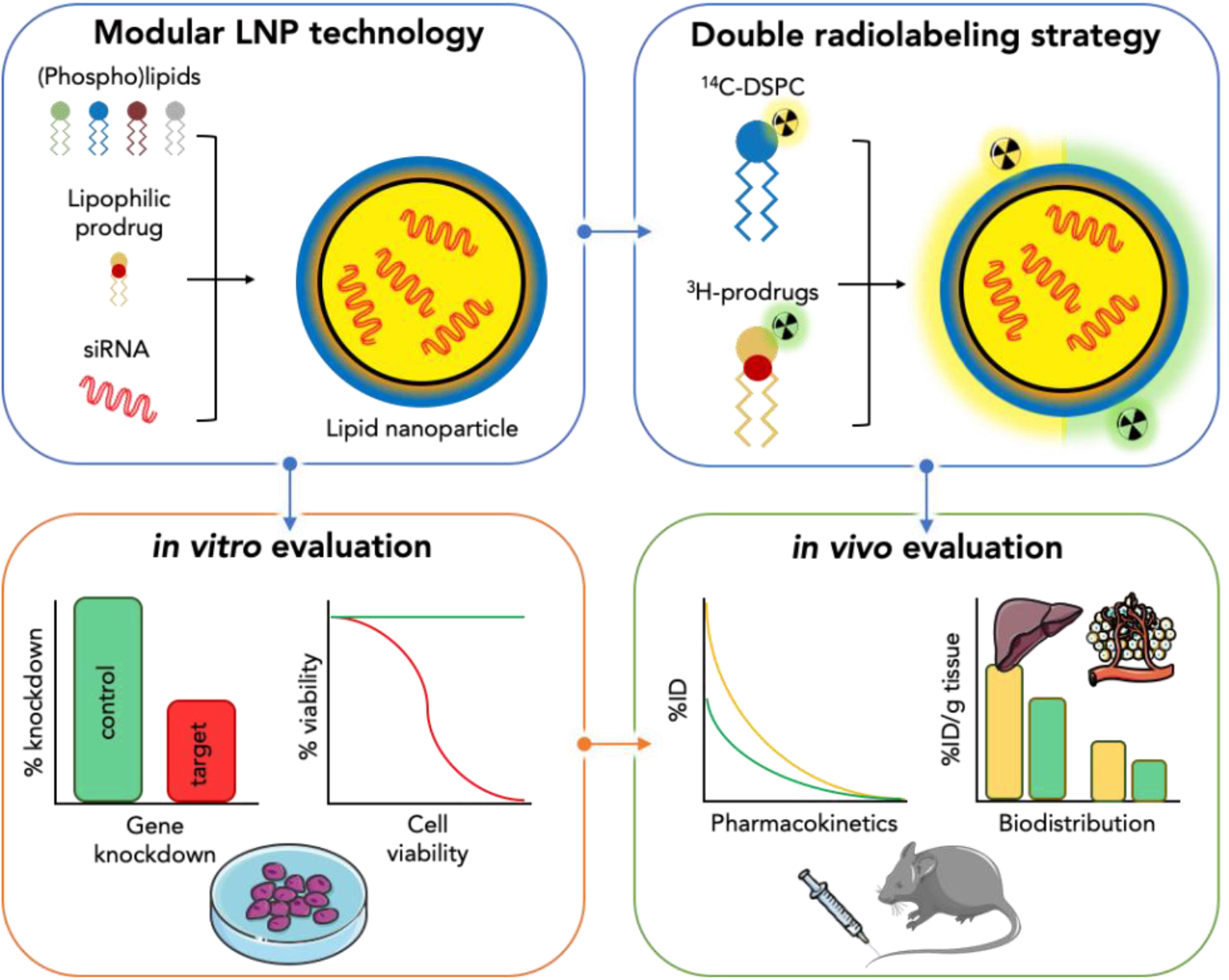

## 1. INTRODUCTION

Treating diseases by silencing pathological genes, expressing therapeutic proteins or via gene editing is increasingly becoming a clinical reality. Several nucleic acid-based therapeutics such as antisense oligonucleotides, small interfering RNA (siRNA), messenger RNA (mRNA) and plasmid DNA have been approved and/or are in late stage clinical trials^1–4^. As genetic drugs can modulate the expression of virtually any gene, they have the potential to therapeutically regulate targets considered ‘undruggable’ by small molecule drugs. However, successfully employing nucleic acids as therapeutics depends heavily on chemical modifications and (nanocarrier) delivery technology to prevent their degradation in the circulation and facilitate intracellular delivery^5^.

The first siRNA therapeutic, called Onpattro^®^, was recently approved for treatment of hereditary transthyretin-mediated amyloidosis^6,7^, twenty years after the discovery of RNA interference^8^. Onpattro^®^ relies on lipid nanoparticle (LNP) technology for siRNA delivery to hepatocytes where it inhibits production of the disease-causing mutant transthyretin protein^9^. LNPs are generally ~50 nm in diameter and consist of cholesterol, phospholipids, polyethylene glycol-conjugated lipids and ionizable cationic lipids. Ionizable amino lipids with an apparent pKa around 6.4 are a key component as their use enables efficient siRNA encapsulation during LNP production at low pH (≤ 4), ensures LNPs’ neutral surface charge in the circulation at physiological pH, and promotes endosomal escape following target cell internalization^10,11^.

One of LNPs’ major advantages hallmarks their applicability as a modular platform technology which facilitates straightforward development of other nucleic acid therapeutics. The siRNA payload is interchangeable as its physicochemical properties remain similar regardless of its sequence. In addition to developing siRNA drugs for hepatic^12^ and extrahepatic^13,14^ targets, LNP technology has also been employed for the therapeutic application of mRNA^15–18^ and gene editing^19^. Moreover, LNPs’ modular design space can be expanded by drug derivatization approaches whereby small molecule prodrugs are synthesized that stably incorporate in LNPs. Incorporating such prodrugs in nanocarriers reduces differences among various drug molecules’ physicochemical characteristics, pharmacokinetic properties and biodistribution profiles, thereby improving drug encapsulation predictability^20^. In addition, these approaches can reduce drug-induced systemic toxicity and improve therapeutic efficacy^21,22^. Finally, modular prodrug designs provide opportunities to co-encapsulate multiple therapeutic agents in a single nanocarrier.

We have recently developed a prodrug platform technology for efficiently incorporating small molecule therapeutics in LNPs^23,24^. The rationally designed prodrugs consist of an active drug coupled via a biodegradable (ester) linker to a hydrophobic moiety. Following LNPs’ cellular uptake, intracellular enzymes break down the linker releasing the active drug. Using this approach, we previously demonstrated that incorporating prodrug derivatives of the corticosteroid dexamethasone in LNP formulations containing nucleic acids significantly reduced their immunostimulatory effects following parenteral administration^23^.

Here, we demonstrate that the modular LNP platform can be used for combined gene silencing and chemotherapy to induce additive anti-cancer effects. As a proof-of-concept, we co-encapsulated siRNA against the N-terminal domain of the androgen receptor (AR), a prostate cancer driver, and various lipophilic prodrug derivatives of the chemotherapeutic agents docetaxel and cabazitaxel. We show that taxane prodrugs can be stably incorporated in LNP-siRNA systems, that prodrug incorporation does not affect the formulation’s ability to knockdown the AR target gene, and that the combination therapy induces additive therapeutic effects *in vitro*. In addition, we used a double-radiolabeling approach to quantitatively determine the LNP and prodrugs’ pharmacokinetic properties and biodistribution following systemic administration. Finally, we show prodrug accumulation and significant AR target gene knockdown in tumors following systemic LNP administration in a murine xenograft tumor model.

## 2. MATERIAL AND METHODS

### 2.1 Materials

1,2-distearoyl-sn-glycero-3-phosphorylcholine (DSPC) was purchased from Avanti Polar Lipids (Alabaster, AL) and cholesterol was obtained from Sigma-Aldrich (St. Louis, MO). The ionizable cationic lipid (6Z,9Z,28Z,31Z)-heptatriaconta-6,9,28,31-tetraen-19-yl4-(dimethylamino) butanoate (DLin-MC3-DMA) was synthesized as previously described by Jayaraman *et al.*^11^, while (R)-2,3-bis(octadecyloxy)propyl-1-(methoxy polyethylene glycol 2000) carbamate (PEG-DMG) and (R)-2,3-bis(stearyloxy)propyl-1-(methoxy poly(ethylene glycol)2000 carbamate (PEG-DSG) were synthesized as previously reported by Akinc *et al*^25^.

### 2.2 Taxane lipophilic prodrug synthesis

Docetaxel and cabazitaxel lipophilic prodrugs were synthesized according to methods previously described by Chen *et al*^23^.

### 2.3 Small interfering RNA (siRNA)

All modified Dicer-substrate siRNA molecules were obtained from Integrated DNA Technologies (IDT, Coralville, IA). The sense and antisense sequences of a previously developed siRNA against exon 1 of AR mRNA (siARNTD) were 5’-ccAuGcAACUCcUuCaGcAACAGdcdA-3’ and 5’-UGcUGUUGcUgAaGGAGUUGCAuGgug-3’^26^. An siRNA against firefly luciferase (siLuc) was used as a negative control (sense, 5’-cuuAcGcuGAGuAcuucGAdTsdT-3’; antisense, 5’-UCGAAGuACUcAGCGuAAGdTsdT-3’)^13^. Lower case letters indicate 2’-O-methyl modification and ‘s’ indicates phosphorothioate linkages between the 3’-deoxythymidine overhangs.

### 2.4 Production of lipid nanoparticles containing siRNA and lipophilic taxane prodrugs

LNPs containing siRNA and lipophilic prodrugs were prepared by rapid mixing through a T-junction mixer as previously described^10,23,27,28^. Briefly, DLin-MC3-DMA, DSPC, cholesterol, PEG-DMG or PEG-DSG and lipophilic prodrugs were dissolved in ethanol at appropriate ratios (Supplementary Table 1) to a final concentration of 10 mM total lipid. Nucleic acids were dissolved in 25 mM sodium acetate buffer at pH 4.0 to obtain a final mixture with a defined nucleic acid to lipid weight/μmol ratio of 0.0278 (N/P 6). The organic and aqueous solutions were mixed at a flow ratio of 1:3 (v:v) and a total flow rate of 28 mL/min. The resulting mixtures were dialyzed against a 1000-fold volume of phosphate buffered saline (PBS) pH 7.4 overnight and sterile filtered (0.2 μm).

**Table 1.**
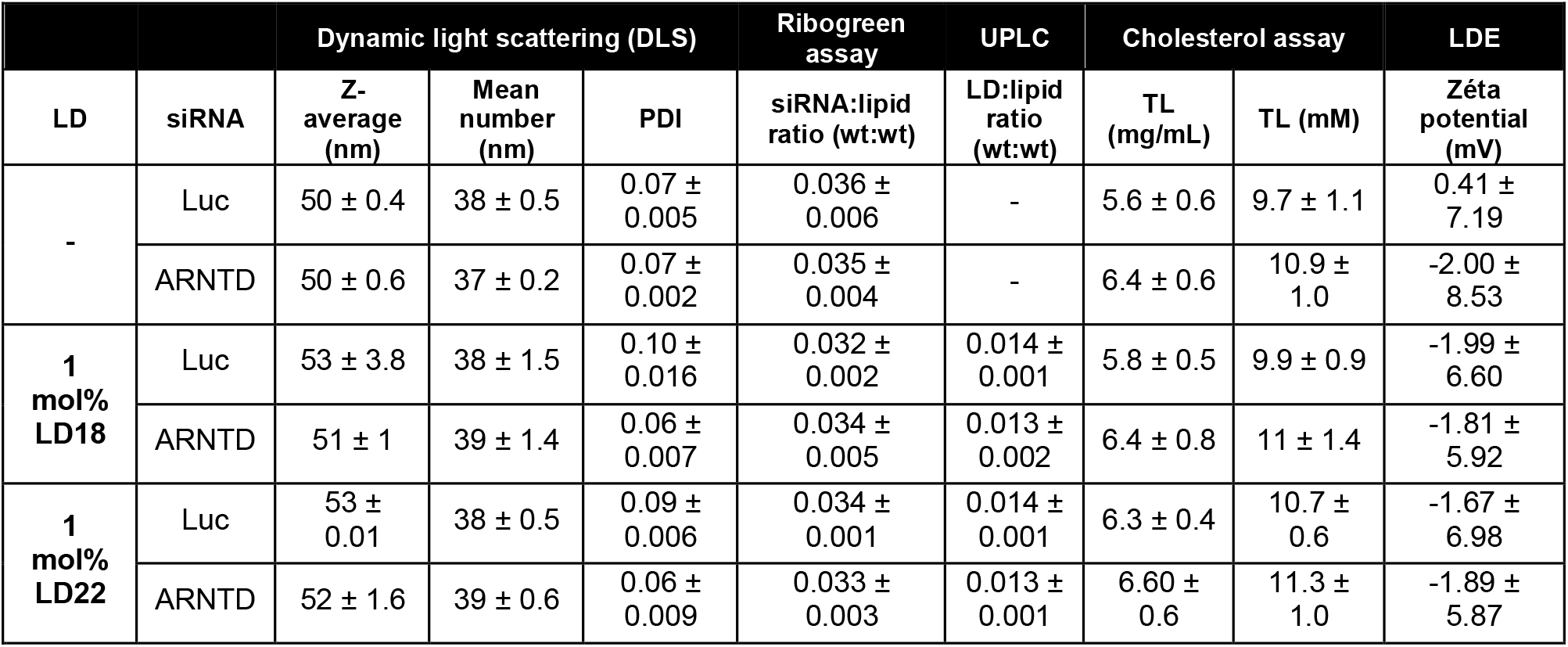
Physicochemical analysis of lipid nanoparticles used for *in vitro* studies. Data are presented as mean ± SD of three representative formulation batches. LNP, lipid nanoparticle; LD, lipophilic docetaxel prodrug; LDE, Laser Doppler Electrophoresis; PDI, polydispersity index; TL, total lipid; wt, weight.

### 2.5 Lipid nanoparticles’ physicochemical analysis

Particle size was determined by dynamic light scattering and zèta potential by laser Doppler electrophoresis using a Malvern Zetasizer NanoZS (Malvern Instruments, Worcestershire, UK). Lipid concentrations were determined by measuring the LNPs’ cholesterol content (Cholesterol E Assay, Wako Chemicals, Richmond, VA) or their phospholipid content (Phospholipids C Assay, Wako Chemicals). The Quant-iT RiboGreen RNA Assay was used to determine siRNA encapsulation and concentration in LNPs according to manufacturer's protocols (ThermoFisher, Waltham, MA). LNP morphology was visualized by cryogenic transmission electron microscopy (cryo-TEM) as previously described^29^. In short, 2-5 μL of concentrated LNPs (20-25 mg/mL total lipid) were added to Lacey-formvar copper grids, and plunge-frozen using a FEI Mark IV Vitrobot (FEI, Hillsboro, OR). Grids were stored in liquid nitrogen until imaged. Grids were moved into a Gatan transfer station pre-equilibrated to at least −180°C prior to add grids to the cryogenic grid holder. A FEI LaB6 G2 TEM (FEI, Hillsboro, OR) operating at 200 kV under low-dose conditions was used to image all samples. A bottom-mount FEI Eagle 4K CCD camera was used to capture all images at a 47-55,000x magnification with a nominal under-focus of 1-2 μm to enhance contrast. Sample preparation and imaging was performed at the University of British Columbia Bioimaging Facility (Vancouver, BC).

### 2.6 *In vitro* degradation of prodrugs

To determine the prodrugs’ biodegradability, 1 mg/mL LNPs containing siLuc and 10 mol% docetaxel prodrugs LD18 or LD22 were incubated in CD1 mouse plasma (Cedarlane, Burlington, Ontario) for 24 hours at 37 °C. Post incubation, four volumes of chloroform/methanol (2:1) were added and vortex mixed. Samples were centrifuged at 13,000g for 5 minutes and the upper phase was discarded. The remaining organic phase was dried down under vacuum and the resulting lipid extract was dissolved in methanol/isopropanol (1:1). Quantity of parent taxane prodrug was determined by ultra-high-performance liquid chromatography (UPLC) on a Waters Acquity H-Class UPLC System equipped with a BEH C18 column (1.7 μm, 2.1 × 100 mm) and a photodiode array detector. Separation was achieved at a flow rate of 0.5 mL/min, with a linear gradient of mobile phases A:B from 30:70 to 100:0 (v:v) over 3 minutes followed by an isocratic hold at 100:0 for an additional 3 minutes. Column temperature was maintained at 55 °C. Mobile phase A consist of equal parts methanol and acetonitrile while mobile phase B is water. The absorbance at 230 nm was measured and the analyte concentration was determined using calibration curves.

### 2.7 Cell lines

The human prostate cancer cell lines 22Rv1 (CRL-2505), LNCaP clone FGC (CRL-1740), PC3 (CRL-1435) and VCaP (CRL-2876) were obtained from the American Type Culture Collection (Manassas, VA). 22Rv1 and LNCaP cells were maintained in RPMI-1640 medium supplemented with 10% (v:v) fetal bovine serum (FBS). PC3 and VCaP cells were maintained in Dulbecco's Modified Eagle Medium (DMEM) supplemented with 10% FBS. All cell lines were kept in culture at 37°C in a humidified atmosphere containing 5% CO2. Cell culture medium and reagents were obtained from Gibco (Thermo Fisher Scientific, Valencia, CA). Cells were propagated to a maximum of 20 passages.

### 2.8 Gene silencing experiments *in vitro*

Cells were seeded at 100,000 – 200,000 cells/well in a 6-well plate and left to adhere overnight in culture medium. The following day, medium was replaced with regular culture medium containing LNP-sRNA at a dose corresponding to 0.05 or 0.1 μg/mL siRNA. After 24- or 48-hours incubation, medium was aspirated and cells were washed three times with cold PBS followed by addition of lysis buffer. Cellular mRNA levels of β-actin and AR variant 7 (AR-V7) were determined as previously described^26^. Briefly, cellular RNA was isolated using an Invitrogen PureLink RNA Mini Kit (Thermo Fisher Scientific) and used as template to synthesize cDNA using a High-Capacity cDNA Reverse Transcription Kit (Applied Biosystems, Foster City, CA). Quantitative PCR (qPCR) was performed using the synthesized cDNA template, TaqMan Fast Advanced Master Mix (Applied Biosystems) and primer-probe assays (IDT) on a Step One Plus Real-Time PCR System (Applied Biosystems). The AR-V7 primer-probe assay included: primer 1, 5’-TTGTCCATCTTGTCGTCTTCG-3’; primer 2, 5’-CAATTGCCAACCCGGAATTT-3’; probe, 5’-TGAAGCAGGGATGACTCTGGGAGA-3’. The β-actin primer-probe assay, used as a household gene for standardizing AR-V7 gene expression levels, included: primer 1, 5’-CCTTGCACATGCCGGAG-3’; primer 2, 5’-ACAGAGCCTCGCCTTTG-3’; probe, 5’-TCATCCATGGTGAGCTGGCGG-3’. Gene probes had a double-quenched design, comprising a 5’ FAM fluorophore, an internal ZEN (IDT) quencher, and a 3’ Iowa Black FQ (IBFQ) quencher. For relative gene expression, AR-V7 mRNA levels were normalized to β-actin mRNA levels using the delta cycle threshold (ΔC^T^) method and the following formula: 2^−ΔCT^ where ΔC^T^ = AR-V7 C^T^ – β-actin C^T^.

### 2.9 Cell viability experiments

The therapeutic effect of free taxane (pro)drugs and LNP formulations was determined by the [3-(4,5-dimethylthiazol-2-yl)-5-(3-carboxymethoxyphenyl)-2-(4-sulfophenyl)-2H-tetrazolium (MTS)-based CellTiter 96 AQueous One Solution Cell Proliferation Assay (Promega) or the resazurin-based PrestoBlue assay (ThermoFisher). For both assays, 10,000 – 15,000 cells/well (22Rv1, LNCaP and VCaP) or 1,500 cells/well (PC3) were seeded in 96-well plates and left to adhere overnight in regular culture medium. The following day, medium was aspirated and replaced with regular culture medium containing increasing concentrations of taxane (pro)drugs or LNP formulations. After 72 or 96 hours, medium containing treatment was aspirated and replaced by medium containing MTS or PrestoBlue solution. Following 0.5 – 4-hour incubation at 37°C, absorbance was measured at 490 nm (MTS) or fluorescence at 530/570 nm (PrestoBlue^®^). Percentage of cell viability was determined by comparing to untreated and DMSO controls.

### 2.10 Pharmacokinetic and biodistribution studies in mice

6-8 weeks old female CD1 mice (Charles River Laboratories, Wilmington) were injected intravenously with radiolabeled LNP-siRNA containing ^14^C-DSPC or ^3^H-CHE and 10 mol% (^3^H-radiolabeled) lipophilic docetaxel prodrugs LD18 or LD22, corresponding to a dose of 2.5 or 4 mg/kg siRNA and ~17 or ~30 mg/kg prodrug. At various timepoints post-injection, mice were anesthetized intraperitoneally with a terminal dose of ketamine-xylazine prior to blood sample collection in EDTA tubes via cardiac puncture. Animals were subsequently euthanized by cervical dislocation followed by tissue collection. Tissues were homogenized in PBS using Fastprep tubes and a Fastprep-24 (MP Biomedical, Santa Ana, CA). Tissue homogenates and blood were subsequently subjected to a digestion and decolorization protocol as previously described^30^. Radioactivity of homogenates, blood samples and LNPs was measured using a Beckman Coulter LS 6500 liquid scintillation counter (Mississauga, Ontario, Canada). The percent recovery in blood was calculated based on a blood volume of 70 ml/kg body weight^30^. Blood and tissue radioactivity are expressed as percent injected dose (%ID) or percent injected dose per gram (%ID/g) of tissue, respectively. All procedures were approved by the Animal Care Committee at the University of British Columbia and were performed in accordance with guidelines established by the Canadian Council on Animal Care.

### 2.11 Biodistribution study in tumor-bearing mice

6-10 weeks old male NRG (NOD-*Rag1_null_ IL2rg_null_*) mice were inoculated subcutaneously on the right flank with 2×10^6^ 22Rv1 cells suspended in 50% Matrigel. Tumor growth was monitored by determining tumor volumes using a digital caliper. The tumor volume *V* (in mm^3^) was calculated using the formula *V = (π/6)LS2* where *L* is the largest and *S* is the smallest superficial diameter. When tumors reached a size of ~250 mm^3^, mice were injected intravenously with LNP-siRNA containing ^14^C-DSPC and 10 mol% ^3^H-radiolabeled docetaxel prodrugs LD18 or LD22, corresponding to a dose of 5 mg/kg siRNA and ~30 mg/kg lipophilic prodrug. At 24 hours post injection, mice were anesthetized with CO_2_ followed by blood collection in EDTA tubes via cardiac puncture. Plasma was separated from whole blood by centrifugation for analysis. Animals were subsequently euthanized followed by tissue and tumor collection. To determine radioactivity, tissue homogenates and plasma were processed as described in Section 2.10. Plasma and tissue radioactivity are expressed as percent injected dose (%ID) or percent injected dose per gram (%ID/g) of tissue, respectively. To determine AR mRNA levels in tumors, total RNA was isolated from tumor homogenates using the TRIzol (Invitrogen) protocol according to the manufacturer’s instructions as previously described^26^. Tumor mRNA levels of β-actin, AR and AR-V7 were determined as described in Section 2.8. All procedures were approved by the Animal Care Committee at the University of British Columbia and BC Cancer Agency and were performed in accordance with guidelines established by the Canadian Council on Animal Care.

### 2.12 Statistical analysis

Curve fitting and statistical analyses were performed with GraphPad Prism 8 software. Pharmacokinetic parameters were analyzed based on a one compartmental analysis of plasma after intravenous bolus using PKSolver^31^.

## 3 RESULTS AND DISCUSSION

### 3.1 Formulating and analyzing lipid nanoparticles containing siRNA and taxane prodrugs

Incorporating lipophilic prodrugs in LNP-siRNA systems has to meet several requirements. First, prodrug incorporation must not disrupt LNP formation and siRNA entrapment during the formulation process. Second, it should not significantly change LNP stability or its physicochemical parameters. Third, the prodrug and siRNA payload’s have to remain their therapeutic activity. To investigate which taxane prodrug is best suited for LNP incorporation, four lipophilic docetaxel derivatives (LD10, LD18, LD22, LD23) and two cabazitaxel derivatives (LD27, LD28) were synthesized by conjugating various hydrocarbon moieties via a biodegradable succinate linker to the parent compounds’ hydroxyl groups (**Figure 1)**^23^.

**Figure 1.**
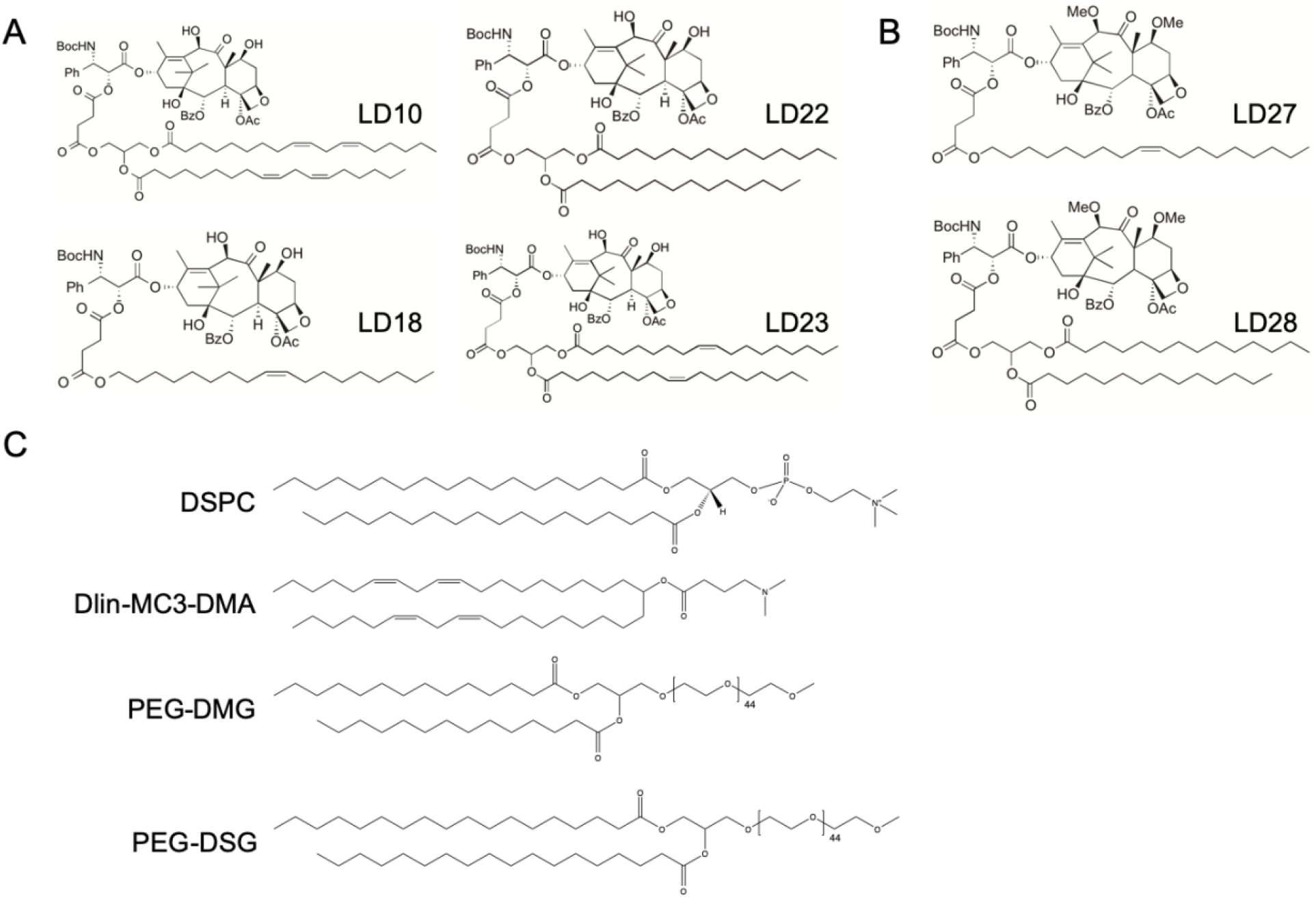
Overview of lipophilic taxane prodrugs and lipids. Structures of (**A**) docetaxel and (**B**) cabazitaxel prodrug derivatives. (**C**) Structures of lipids used for LNP formation. DSPC,1,2-distearoyl-sn-glycero-3-phosphorylcholine; DLin-MC3-DMA, (6Z,9Z,28Z,31Z)-Heptatriaconta-6,9,28,31-tetraen-19-yl4-(dimethylamino) butanoate; PEG-DMG, (R)-2,3-bis(octadecyloxy)propyl-1-(methoxy polyethylene glycol 2000) carbamate; PEG-DSG, (R)-2,3-bis(stearyloxy)propyl-1-(methoxy poly(ethylene glycol)2000 carbamate.

The synthesized lipophilic taxane prodrugs and siRNA were incorporated in LNPs by a rapid mixing procedure using a T-junction device as previously described^10,23,27,28^. The lipid compositions, based on the optimized LNP formulation used for Onpattro^®9,11^, are listed in **Supplementary Table 1.**

As shown in **Table 1**, LNP-siRNA formulations containing 1 mol% docetaxel prodrugs LD18 or LD22 used for *in vitro* studies, had particle diameters of approximately 50 nm (Z-average) with polydispersity indices (PDI) ≤ 0.1 and nearly neutral zèta potentials. LNP formulations containing 10 mol% LD18 or LD22 used for *in vitro* and *in vivo* (**Supplementary Table 2-5**) studies had comparable physicochemical characteristics. This indicates that prodrug incorporation up to 10 mol% did not significantly affect LNP size, homogeneity or zèta potential. These results were corroborated by cryogenic transmission electron microscopy (cryoTEM), which demonstrated that LNPs containing 10 mol% LD18 or LD22 displayed similar morphology and size when compared to control formulations without prodrugs (**Figure 2**). Efficient prodrug and siRNA entrapment (routinely > 90%) were observed for all formulations containing 0.1 - 10 mol% taxane prodrugs.

**Figure 2.**
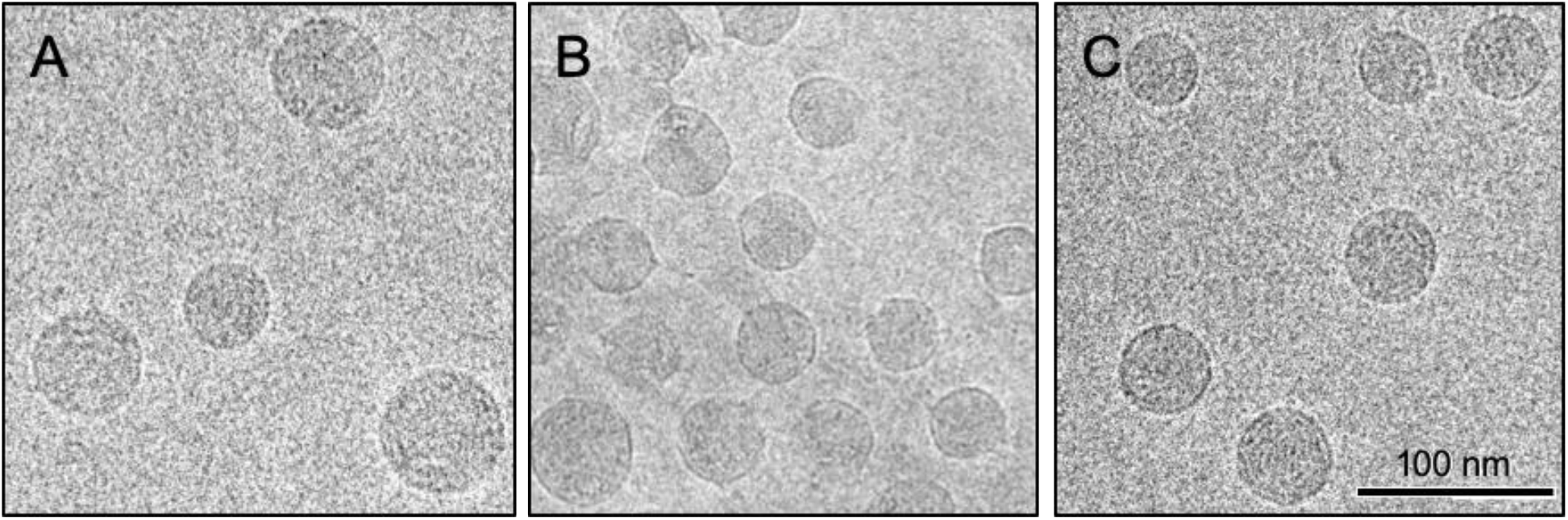
Cryogenic transmission electron micrographs of lipid nanoparticles containing siRNA and lipophilic docetaxel prodrugs. (**A**) Lipid nanoparticles containing luciferase siRNA and 10 mol% docetaxel prodrugs (**B**) LD18 or (**C**) LD22 were visualized using cryogenic transmission electron microscopy operating at 200 kV.

The taxane prodrugs’ cytotoxic activity requires their release from the LNP-siRNA systems and subsequent liberation of the active compounds mediated by esterases. To assess this, LNP-siRNA containing 10 mol% docetaxel prodrugs LD18 or LD22 were incubated with esterase-rich mouse plasma and the amount of parent prodrug was determined. After 24-hour incubation, approximately 75% and 50% of LNP-associated LD18 and LD22, respectively, were degraded (**Figure 3A**). The variation in degradation could be caused by differences in prodrug lipophilicity and LNP incorporation. It is possible that the more lipophilic LD22 containing two saturated C^14^ lipid chains incorporates more stably in the hydrophobic LNP core during production and is therefore less accessible to esterases. Similar results were observed with incorporating lipophilic dexamethasone prodrugs in LNP-siRNA^23^.

**Figure 3.**
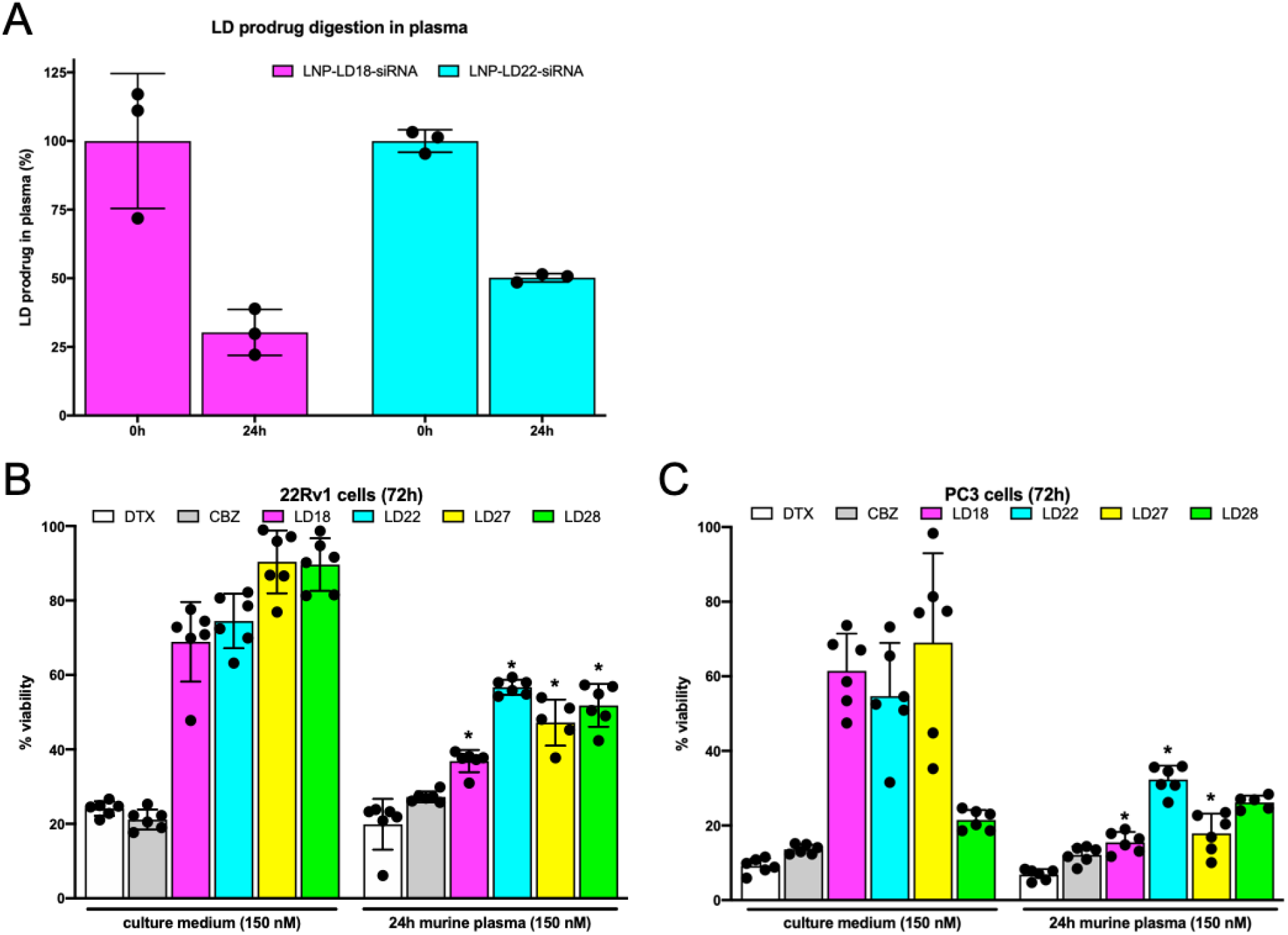
Esterase-mediated hydrolytic activation of lipophilic taxane prodrugs. (**A**) Docetaxel prodrug digestion in plasma. LNPs containing luciferase siRNA and 10 mol% docetaxel prodrug LD18 or LD22 were incubated with CD1 mouse plasma for 24 hours. Prodrug concentration in plasma samples was determined by UPLC analysis. Reduction in parent prodrug amount indicates digestion in plasma. Data represents mean ± SD (n=3) of one representative experiment. **(B, C)** Incubating taxane prodrugs with murine plasma liberates active cytotoxic parent compounds. Docetaxel ( DTX) prodrugs (LD10, LD18, LD22, LD23) and cabazitaxel (CBZ) prodrugs (LD27, LD28) were incubated in culture medium or CD1 mouse plasma at 37°C for 24 hours prior to 72-hour (**B**) 22Rv1 or (**C**) PC3 cell treatment. Cytotoxicity was determined by MTS cell viability assay. Data are presented as mean ± SD of one representative experiment (n=6) and analyzed by two-way ANOVA with Tukey’s post-test. * indicates p-value <0.005 plasma incubation vs. culture medium incubation.

To determine if liberated taxane compounds remain their functionality and induce cytotoxicity, prodrugs were first incubated with mouse plasma followed by exposure to prostate cancer cells. As shown in **Figure 3B** and **C,** taxane prodrugs significantly reduce 22Rv1 and PC3 cell viability when pre-incubated with murine plasma as compared to prodrugs pre-incubated in regular culture medium. This is not observed for unmodified docetaxel and cabazitaxel, which are equally potent regardless of culture medium or mouse plasma pre-incubation. This indicates that derivatizing taxane chemotherapeutics for stable LNP incorporation likely reduces their toxicity, while their cytotoxic activity is restored by liberation of the active compound in an esterase-rich environment.

### 3.2 LNPs containing androgen receptor siRNA and taxane prodrugs induce additive therapeutic effects *in vitro*

We next assessed LNPs’ therapeutic efficacy *in vitro* by determining their ability to induce target gene knockdown and inhibit cell viability using various genetically distinct prostate cancer cell lines. In 22Rv1 cells, which express wild type AR and the frequently occurring splice variant AR-V7, LNPs containing AR N-terminal domain (ARNTD) siRNA significantly silenced AR-V7 expression at doses as low as 0.01 μg/mL siRNA when compared to control LNP formulation containing luciferase siRNA (**Supplementary Figure 1A-C**). To evaluate if taxane prodrug incorporation affects LNP-siRNA’s gene knockdown activity, 22Rv1 and LNCaP cells were incubated with LNPs containing 1 mol% taxane prodrugs and ARNTD siRNA, followed by determining AR-V7 expression levels. As shown in **Figure 4A** and **B**, LNPs containing ARNTD siRNA and docetaxel or cabazitaxel prodrugs induce significant AR-V7 knockdown when compared to control formulations containing prodrugs and luciferase siRNA at a dose of 0.05 μg/mL siRNA. These results show that prodrug incorporation does not affect LNP-siRNA’s gene silencing ability.

**Figure 4.**
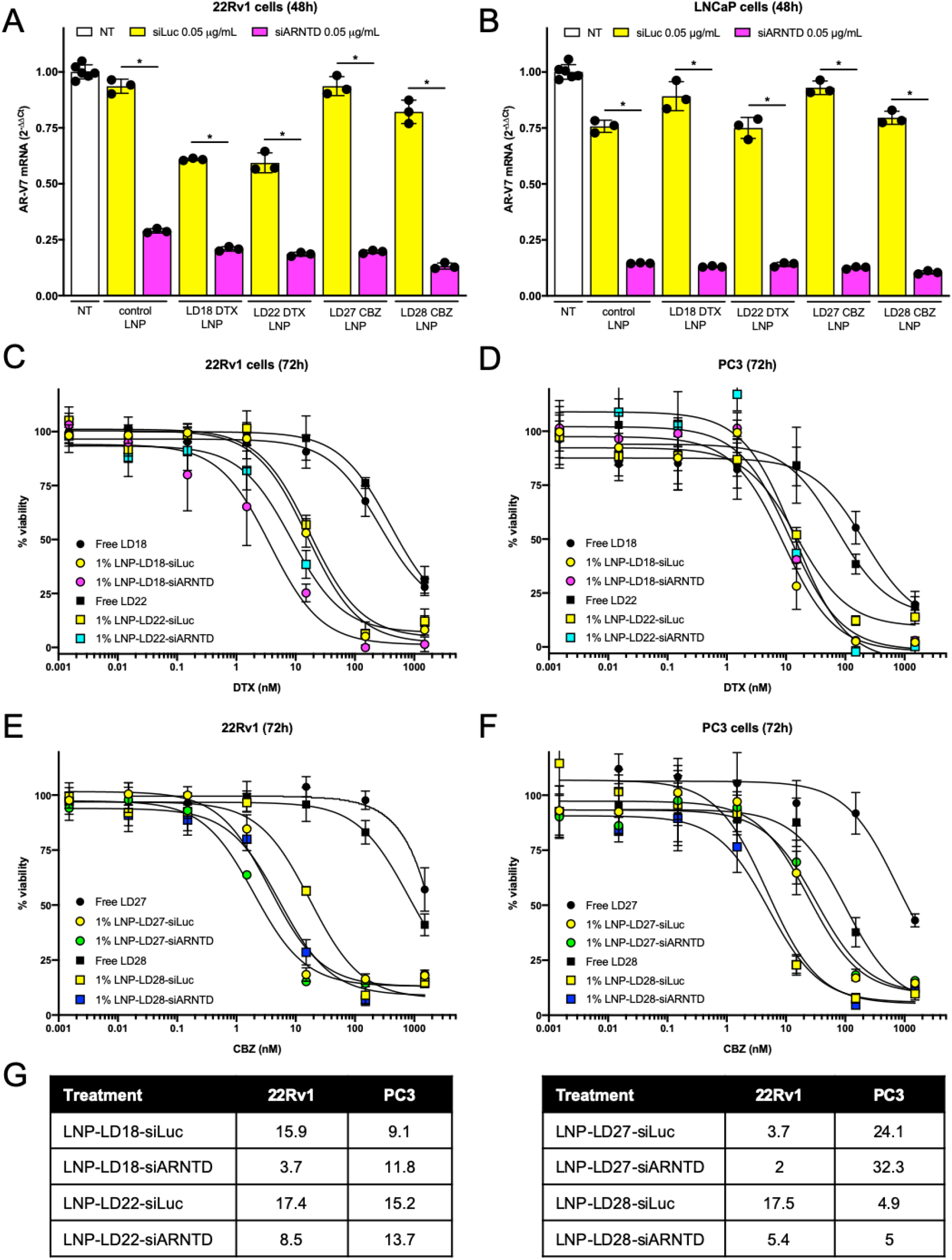
Lipid nanoparticles containing taxane prodrugs and androgen receptor siRNA induce target gene knockdown and inhibit cell viability. (**A**) 22Rv1 and (**B**) LNCaP cells were exposed for 48 hours to LNPs containing 1 mol% docetaxel (DTX; LD18, LD22) or cabazitaxel (CBZ; LD27, LD28) prodrug and siRNA against luciferase (siLuc) or androgen receptor (AR) N-terminal domain (siARNTD) at a dose of 0.05 μg/mL. RT-qPCR was used to determine mRNA levels. AR variant 7 (AR-V7) target gene expression change was calculated relative to β-actin control. Data represent mean ± SD (n=3) of one representative experiment and analyzed by one-way ANOVA with Tukey’s post-test. * indicates p-value <0.0001 of LNP formulations containing siARNTD vs. siLuc control. (**C, E**) 22Rv1 and (**D, F**) PC3 cells were exposed for 72 hours to LNPs containing 1 mol% taxane prodrugs and siLuc or siARNTD. Cell viability was determined by MTS assay. Data are presented as mean ± SD of one representative experiment (n=6). (**G**) IC50 values (nM prodrug) derived from data in panels **C-F** determined by Graphpad Prism 8.

Exposing 22Rv1 cells for 96 hours to LNPs containing ARNTD siRNA at a dose of 0.05 μg/mL siRNA results in significant cytotoxicity, while no decrease in cell viability is observed for LNPs containing luciferase siRNA (**Supplementary Figure 1D**). Importantly, exposing PC3 cells, which do not express AR or AR-V7, to the same siRNA dose does not result in cell viability inhibition for LNPs containing either ARNTD or luciferase siRNA (**Supplementary Figure 1D**). This indicates that the cell viability inhibition is a result of specific AR/AR-V7 knockdown induced by LNPs containing ARNTD siRNA.

To determine if AR/AR-V7 knockdown and taxane-induced cytotoxicity would induce additive therapeutic effects, we first exposed prostate cancer cells to increasing concentrations of docetaxel or cabazitaxel combined with LNPs containing ARNTD siRNA at a constant dose of 0.05 μg/mL. As shown in **Supplementary Figure 2**, additive cytotoxic effects are observed in AR/AR-V7-expressing 22Rv1 and VCaP cells, while this is not the case for PC3 cells. Encouraged by these results, we proceeded to expose 22Rv1 and PC3 cells to increasing concentrations of LNPs containing 1 mol% lipophilic taxane prodrugs and ARNTD siRNA (**Figure 4**). In both cell lines, LNPs containing taxane prodrugs and ARNTD or luciferase siRNA induce dose-dependent cell viability inhibition. However, in 22Rv1 cells, LNPs with ARNTD siRNA are considerably more potent than control formulations with luciferase siRNA. For example, the IC50 value of LNPs containing ARNTD and docetaxel prodrug LD18 is 3.7 nM (prodrug concentration), while it is 15.9 nM for the control formulation. For formulations containing docetaxel prodrug LD22, these values are 8.5 nM for the LNPs containing ARTND siRNA versus 17.4 nM for the control formulation (**Figure 4C, G**). In contrast, in PC3 cells, the IC50 values are comparable for LNPs containing taxane prodrugs and ARNTD or luciferase siRNA (**Figure 4D, G**). Similar results were observed for formulations containing cabazitaxel prodrugs (**Figure 4E** and **F**) and LNPs containing 0.1 mol% taxane prodrugs (**Supplementary Figure 3**). Of note, the activity of free docetaxel and cabazitaxel prodrugs in both cell lines is limited, while prodrug incorporation in LNPs containing luciferase siRNA results in dose-dependent cell viability inhibition. This indicates that taxane prodrug derivatization impairs their cell uptake, possibly reducing taxane-mediated toxicity.

### 3.3 Taxane prodrugs remain associated with lipid nanoparticles following systemic administration

Using prostate cancer xenograft mouse models, it has previously been shown that LNP-siRNA can accumulate in tumors and induce therapeutic effects following systemic administration^14,26,32^. LNPs’ tumor accumulation largely depends on their ability to stay intact and circulate sufficiently long to reach tumor cells. To determine if taxane prodrug incorporation affects LNP circulation time, we formulated radiolabeled LNP-siRNA containing ^3^H-labeled taxane prodrugs and conducted pharmacokinetic studies in CD1 mice following systemic administration. This double radiolabel approach allowed us to quantitively track both the LNP and the taxane prodrug payload, making it possible to determine if the prodrug remains associated with the LNP over time following injection.

We first determined control LNP-siRNA’s circulation time, using ^14^C-labeled DSPC and ^3^H-labeled cholesteryl hexadecyl ether (CHE) in the formulation. As shown in **Figure 5A**, percentage of injected dose quantification at various timepoints provided identical outcomes when determined by ^14^C-labeled DSPC (T_1/2_ 4.63 hours) or ^3^H-CHE (T_1/2_ 4.55 hours). This demonstrates that incorporating ^14^C-labeled DSPC in LNPs is a reliable strategy which, combined with ^3^H-labeled prodrugs, can be used to determine both the nanocarrier and the prodrug payload’s pharmacokinetic parameters following intravenous injection in mice.

**Figure 5.**
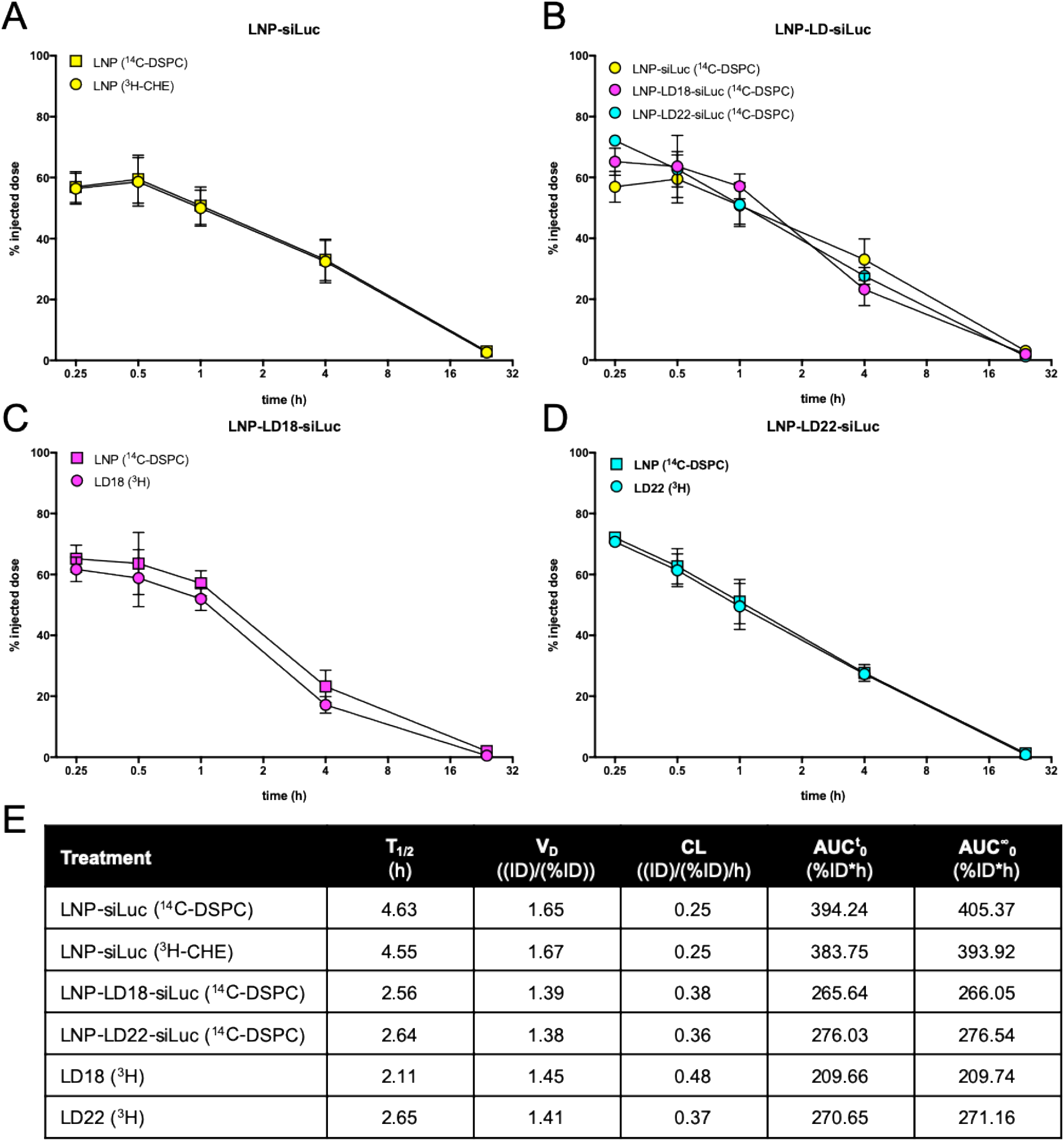
Pharmacokinetic parameters of double radiolabeled lipid nanoparticles following systemic administration in mice. LNP-siRNA containing 10 mol% docetaxel prodrug LD18 or LD22 had a double radiolabel to track both the LNP as well as the prodrugs in the circulation after injection. CD1 mice were injected intravenously with LNPs at a dose of 4 mg/kg siRNA, corresponding to ± 30 mg/kg docetaxel prodrug. Blood samples were collected via heart puncture at various timepoints and radiolabels were quantifiied by liquid scintillation counting. (**A**) Circulation time of control LNPs containing siLuc determined by 3H-CHE and 14C-DSPC. (**B**) Comparison of LNP circulation times as determined by 14C-DSPC. (**C**) Circulation time of LNP-LD18-siLuc determined by 14C-DSPC (LNP) and 3H-LD18 (prodrug). (**D**) Circulation time of LNP-LD22-siLuc determined by 14C-DSPC (LNP) and 3H-LD22 (prodrug). (**E**) Data are presented as mean ± SD of one experiment (n=3-4 animals per timepoint). Pharmacokinetic parameters of LNP formulations determined by PKSolver. T_1/2_, elimination half-life; VD, volume of distribution; CL, clearance; AUC, area under the curve.

Although control LNP-siRNA and formulations containing LD18 or LD22 prodrugs show comparable pharmacokinetic profiles as determined by ^14^C-DSPC (**Figure 5B**), prodrug incorporation affects LNP circulation time to some extent. With a T_1/2_ of 2.6 hours for LNPs containing prodrugs, it is slightly shorter than control LNP-siRNA (**Figure 5E**). It is possible that the lower cholesterol amount present in the formulation containing prodrugs affects the LNP’s opsonization and subsequent blood clearance^33^.

Importantly, the circulation time of the ^3^H-labeled LD18 (T_1/2_ 2.11 hours) and LD22 (T_1/2_ 2.65 hours) prodrugs is comparable to that of their respective LNP-siRNA (**Figure 4C, D**), indicating that the prodrugs remain associated with the LNPs in the circulation. This suggests that LNPs have the potential to simultaneously deliver their siRNA and prodrug payload to target cells following systemic administration, which is important for their therapeutic effect.

Of note, LNP-siRNA’s pharmacokinetic parameters deviate somewhat from results observed in a previous study (**Supplementary Figure 4**). For example, while LNP-siRNA’s T_1/2_ (4.55 hours) is longer than we previously observed (T_1/2_ 3.04 hours), the T_1/2_ for LNP-siRNA containing LD18 (2.56 hours) or LD22 (2.64 hours) is slightly shorter than in the previous study (3.06 hours and 3.72 hours, respectively). Although the reasons for the observed variations is unclear, differences in injected dose, radiolabel and quantification method could have influenced the results.

### 3.4 Lipid nanoparticles deliver siRNA and taxane prodrugs to tumors

To determine to what extent LNP-siRNA and associated taxane prodrugs accumulate in tumors following systemic administration, a biodistribution study was conducted using a 22Rv1 subcutaneous tumor xenograft model. Similar as for the pharmacokinetic study, ^14^C-radiolabeled LNP-siRNA containing ^3^H-labeled prodrugs were used to determine the individual tissue biodistribution of both the LNPs and the prodrug payload.

**Figure 6A-C** show the tissue distribution of the control formulation and LNP-siRNA containing docetaxel prodrugs LD18 or LD22, 24 hours following intravenous injection. Despite differences in pharmacokinetic parameters, unmodified and prodrug-containing LNPs show comparable tissue distribution. The highest LNP accumulation is observed in the liver (± 15% injected dose per gram tissue, ID/g tissue). This is expected as LNPs have been designed to transfect hepatocytes^10,11^. LNP accumulation was also observed in the spleen (± 4-5% ID/g tissue), likely due to uptake by phagocytic cells. LNPs accumulated to a comparable degree in the kidneys and to a lesser extent in the lungs and heart. Importantly, for all LNP formulations, approximately 2% ID/g tissue accumulated in the tumor (**Figure 6D**). This is in line with other nanocarriers’ accumulation in solid tumors^34^.

**Figure 6.**
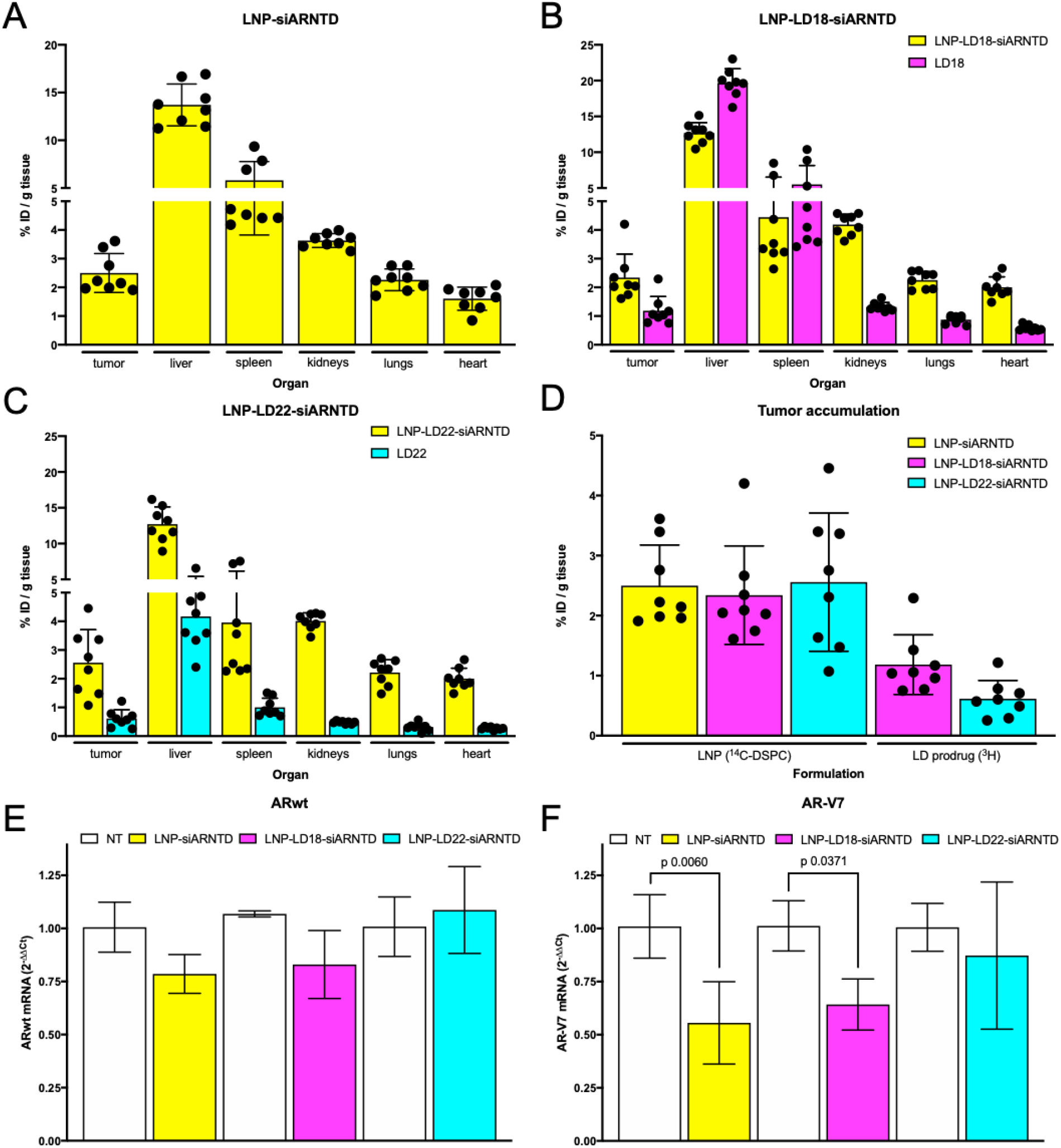
Lipid nanoparticles accumulate in tumors and induce target gene knockdown following systemic administration in mice. NRG mice bearing 22Rv1 tumors were injected intravenously with ^14^C-DSPC-labeled LNP-siRNA containing 10 mol% 3H-labeled docetaxel prodrug LD18 or LD22 at a dose of 5 mg/kg siRNA, corresponding to ± 30 mg/kg docetaxel prodrug. After 24 hours, mice were sacrificed and tissues were collected for radiolabel quantification by liquid scintillation counting. Tissue distribution profiles of (**A**) control LNPs containing ARNTD siRNA (LNP-siARNTD), (**B**) LNPs containing siARNTD and LD18 (LNP-LD18-siARNTD), and (**C**) LNPs containing siARNTD and LD22 (LNP-LD22-siARNTD). (**D**) Tumor accumulation of LNP formulations. Radioactivity is expressed as percent injected dose per gram (%ID/g) of tissue. Data represent mean ± SD (n=8 animals). (**E, F**) RT-qPCR was used to determine tumor mRNA levels. (**E**) AR wild type (ARwt) and (**F**) AR variant 7 (AR-V7) target gene expression change was calculated relative to β-actin control. Data represent mean ± SD (n=8 animals) of one experiment and analyzed by one-way ANOVA with Tukey’s post-test.

Notable differences were observed for LD18 and LD22’s tissue distribution (**Figure 6B, C**). For example, LD18 accumulated in the liver (± 20% ID/g tissue) and spleen (± 5% ID/g tissue), even to a higher extent than the associated LNP. In contrast, LD22’s liver (± 4% ID/g tissue) and spleen (± 1% ID/g tissue) accumulation were considerable lower. As shown in **Figure 6D**, LD18’s overall higher tissue accumulation compared to LD22 was also observed in the tumor (± 1.2% versus 0.8% ID/g tissue, respectively). It is unclear what caused the differences in prodrug tissue accumulation, since the associated LNPs’ tissue accumulation was comparable. Additional experiments with LNPs containing other taxane prodrug derivatives are required to determine how the structure and lipophilicity influence tissue accumulation.

Unmodified LNP-siRNA have previously been shown to be capable of inducing gene silencing in prostate cancer xenograft models^26,32^. To determine if LNPs containing taxane prodrugs and ARNTD siRNA could induce AR/AR-V7 knockdown, RNA was extracted from tumors collected during the biodistribution study and analyzed by RT-qPCR (**Figure 6E, F**). While accumulating in tumors to a comparable extent as the other formulations, LNPs containing LD22 did not induce knockdown of AR or AR-V7. In contrast, control LNPs containing ARNTD siRNA and formulations containing siRNA and LD18 induced a slight reduction in AR mRNA levels, and significantly reduced AR-V7 levels compared to non-treated controls.

Collectively, these results indicate that LNP-siRNA containing lipophilic taxane prodrugs are able to deliver their therapeutic payloads to tumors, resulting in target gene knockdown.

## 4. CONCLUSIONS

LNP nanocarrier technology has enabled the clinical translation of RNA-based gene therapies. At the same time, LNPs’ modularity and drug derivatization methods provide opportunities to develop rational combination strategies. For example, incorporating dexamethasone prodrugs to reduce the immunostimulatory properties of LNPs containing nucleic acids^23^.

Here, we show that incorporating taxane prodrugs in LNP-siRNA systems offers an attractive strategy to induce additive anti-cancer effects via combined gene silencing and chemotherapy. Various lipophilic docetaxel and cabazitaxel derivatives were stably incorporated in LNPs without affecting its physicochemical properties or compromising LNP-siRNA’s gene silencing ability. As a proof-of-concept, combined LNP-siRNA-mediated knockdown of AR, a prostate cancer driver, and taxane prodrug-induced cytotoxicity induced additive therapeutic effects *in vitro*. Finally, we demonstrated that prodrugs remain associated with LNP-siRNA following systemic administration, resulting in prodrug accumulation and target gene knockdown in tumors *in vivo*. Our study indicates that co-encapsulation of siRNA and lipophilic prodrugs into LNPs provides opportunities for rational design of combination therapies.

## ACKNOWLEDGMENTS

The authors are grateful for the dedicated support of dr. Nancy Dos Santos and Nicole Wretham (BC Cancer Agency) during studies with tumor-bearing mice, Dominik Witzigmann (University of British Columbia) for assisting with pharmacokinetic analysis and Joslyn Quick (University of British Columbia) for assisting with tissue processing and RNA analysis. RvdM’s research on LNPs is supported by funding from the European Union’s Horizon 2020 research and innovation program under the Marie Sklodowska-Curie grant agreement No. 660426 and a Veni STW Fellowship (#14385) from the Netherlands Organization for Scientific Research (NWO). The work undertaken by JZ, JAK and PRC is funded by a Foundation Grant (FDN 148469) from the Canadian Institute for Health Research (CIHR). The work of RMS on LNPs has received funding from the European Union’s Horizon 2020 research and innovation program under grant agreement No. 721058.

## CONTRIBUTIONS

RvdM designed all experiments, formulated all LNPs used throughout the study and performed physicochemical analyses, conducted *in vitro* and *in vivo* studies, and wrote the manuscript.

SC co-designed experiments, assisted with LNP production and analysis, characterization studies, (radiolabeled) lipophilic prodrug synthesis, tissue collection and processing, and reviewed the manuscript.

JZ synthesized DLin-MC3-DMA, PEG lipids and (radiolabeled) lipophilic taxane prodrugs for all studies, and reviewed the manuscript.

JAK assisted with LNP production and analysis, conducted cryoTEM imaging, tissue collection and processing, and reviewed the manuscript.

XRSZ assisted with LNP production and analysis, cell viability experiments and qPCR studies. YKT performed injections and tissue collection.

MBB co-designed the biodistribution study and reviewed the manuscript. RMS co-designed experiments and co-wrote the manuscript.

PRC co-designed experiments and co-wrote the manuscript.

YYCT co-designed and supervised all experiments, assisted in tissue collection and co-wrote the manuscript.

