## Supplemental Information for "Modular lipid nanoparticle platform technology for siRNA and lipophilic prodrug delivery"

### SUPPLEMENTARY TABLES

| LNP formulation composition for *in vitro* studies | | | | | |
|---|---|---|---|---|---|
| Formulation | DLin-MC3-DMA | DSPC | cholesterol | PEG-DMG | Lipophilic prodrug |
| Control | 45 | 11 | 42.5 | 1.5 | - |
| 0.1 mol% LD | 45 | 10.9 | 42.5 | 1.5 | 0.1 |
| 1 mol% LD | 45 | 10 | 42.5 | 1.5 | 1 |
| 5 mol% LD | 45 | 10 | 38.5 | 1.5 | 5 |
| 10 mol% LD | 45 | 9 | 34.5 | 1.5 | 10 |
| LNP formulation composition for *in vivo* studies | | | | | |
| Formulation | DLin-MC3-DMA | DSPC | cholesterol | PEG-DSG | Lipophilic prodrug |
| Control | 50 | 10 | 37.5 | 2.5 | - |
| 10 mol% LD | 45 | 8 | 34.5 | 2.5 | 10 |

**Supplementary Table 1. Lipid nanoparticle compositions for *in vitro* and *in vivo* studies.** Lipid compositions in mol%. LD, lipophilic taxane derivative; LNP, lipid nanoparticle; DLin-MC3-DMA, heptatriaconta-6,9,28,31-tetraen-19-yl 4-(dimethylamino)butanoate; DSPC, 1,2-distearoyl-sn-glycero-3-phosphocholine; PEG-DMG, polyethylene glycol-dimyristolglycerol; PEG-DSG, polyethylene glycol-distearoylglycerol.

| | | | Dynamic light scattering (DLS) | | | Ribogreen assay | UPLC | Cholesterol assay | | LDE |
|---|---|---|---|---|---|---|---|---|---|---|
| **LNP** | **LD** | **siRNA** | **Z-average (nm)** | **Mean number (nm)** | **PDI** | **siRNA:lipid ratio (wt:wt)** | **LD:lipid ratio (wt:wt)** | **TL (mg/mL)** | **TL (mM)** | **Zéta potential (mV)** |
| **10 mol%** | LD18 | Luc | 58 ± 0.1 | 44 ± 1.4 | 0.066 ± 0.019 | 0.037 | 0.11 | 4.14 | 6.36 | -2.16 ± 7.78 |
| | LD22 | | 55 ± 1.5 | 39 ± 0.8 | 0.111 ± 0.031 | 0.037 | 0.10 | 4.31 | 6.38 | -3.11 ± 8.38 |

**Supplementary Table 2. Physicochemical analysis of lipid nanoparticles prepared for cryogenic electron microscopy and digestion study *in vitro*.** DLS data are presented as mean ± SD for one formulation batch measured in triplicate. LNP, lipid nanoparticle; LD, lipophilic docetaxel derivative; LDE, Laser Doppler Electrophoresis; PDI, polydispersity index.

| | | | Dynamic light scattering (DLS) | | | Ribogreen assay | UPLC | Cholesterol assay | |
|---|---|---|---|---|---|---|---|---|---|
| **LNP** | **LD** | **siRNA** | **Z-average (nm)** | **Mean number (nm)** | **PDI** | **siRNA:lipid ratio (wt:wt)** | **LD:lipid ratio (wt:wt)** | **TL (mg/mL)** | **TL (mM)** |
| **10 mol%** | - | Luc | 48 ± 0.19 | 33 ± 0.54 | 0.110 ± 0.010 | 0.032 | - | 30.31 | 50.73 |
| | LD18 | Luc | 51 ± 0.19 | 38 ± 0.59 | 0.060 ± 0.006 | 0.027 | 0.17 | 18.57 | 27.85 |
| | LD22 | Luc | 51 ± 0.37 | 38 ± 1.51 | 0.070 ± 0.010 | 0.028 | 0.20 | 23.99 | 34.72 |

**Supplementary Table 3. Physicochemical analysis of lipid nanoparticles for pharmacokinetic study.** DLS data are presented as mean ± SD for one formulation batch measured in triplicate. LNP, lipid nanoparticle; TL, total lipid; LD, lipophilic docetaxel derivative; PDI, polydispersity index.

| | | | Dynamic light scattering (DLS) | | | Ribogreen assay | UPLC | Cholesterol assay | |
|---|---|---|---|---|---|---|---|---|---|
| **LNP** | **LD** | **siRNA** | **Z-average (nm)** | **Mean number (nm)** | **PDI** | **siRNA:lipid ratio (wt:wt)** | **LD:lipid ratio (wt:wt)** | **TL (mg/mL)** | **TL (mM)** |
| **10 mol%** | - | ARNTD | 44 ± 0.28 | 29 ± 0.83 | 0.118 ± 0.008 | 0.033 | - | 36.43 | 59.69 |
| | LD18 | ARNTD | 51 ± 0.48 | 35 ± 2.03 | 0.099 ± 0.021 | 0.031 | 0.17 | 38.31 | 57.47 |
| | LD22 | ARNTD | 48 ± 0.08 | 36 ± 0.92 | 0.072 ± 0.004 | 0.028 | 0.19 | 37.50 | 54.26 |

**Supplementary Table 4. Physicochemical analysis of lipid nanoparticles for biodistribution study.** DLS data are presented as mean ± SD (n=3) for one formulation batch measured in triplicate. LNP, lipid nanoparticle; TL, total lipid; LD, lipophilic docetaxel derivative; PDI, polydispersity index.

### SUPPLEMENTARY FIGURES

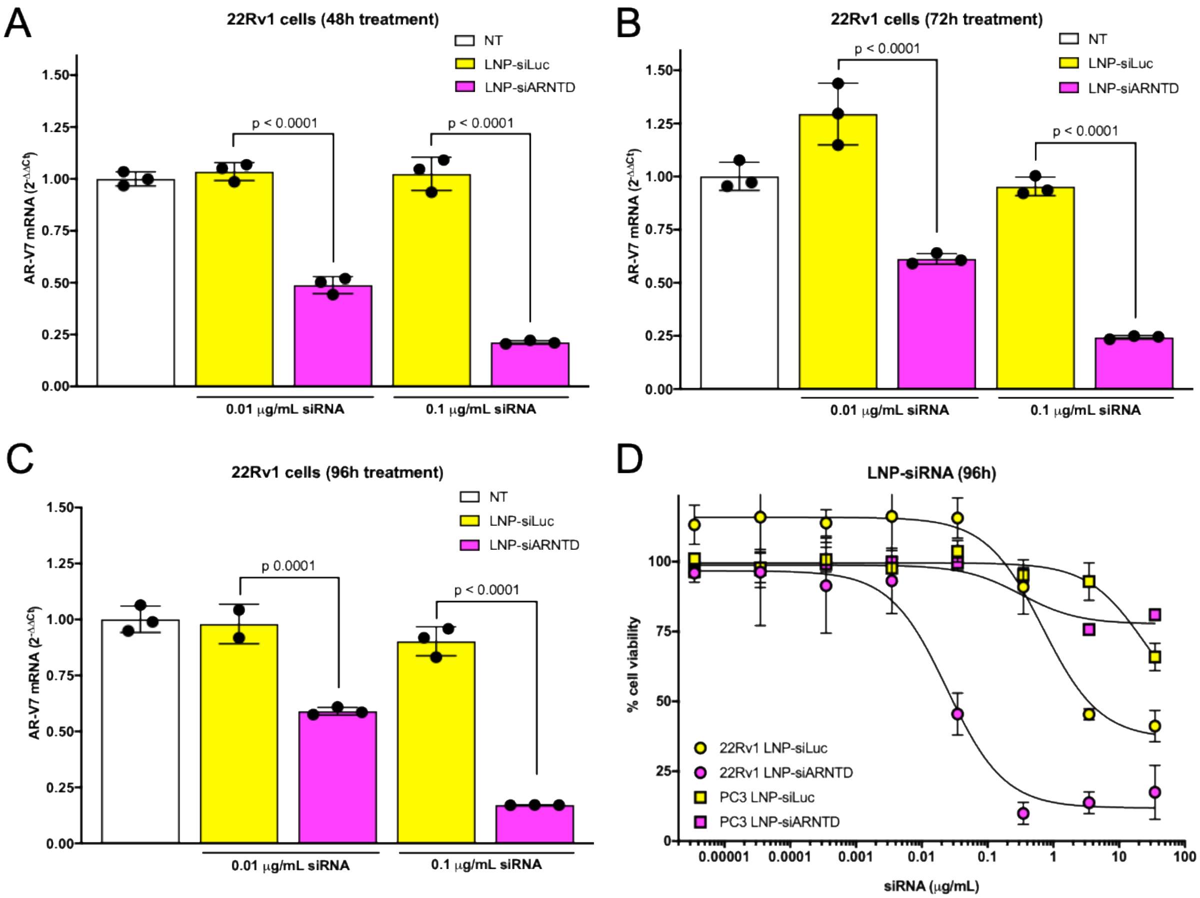


**Supplementary Figure 1. Lipid nanoparticles containing androgen receptor siRNA induce target gene knockdown and inhibit cell viability**. 22Rv1 cells were exposed for **(A)** 24 hours, **(B)** 72 hours or **(C)** 96 hours to LNPs containing androgen receptor N-terminal domain siRNA (LNP-siARNTD, magenta) or luciferase siRNA (LNP-siLuc, yellow) in concentrations corresponding to 0.01 or 0.1 µg/mL siRNA. RT-qPCR was used to determine mRNA levels. AR-V7 target gene expression change was calculated relative to β-actin control. Data are presented as mean ± SD (n=3) of one representative experiment and analyzed by one-way ANOVA with Tukey's post-test. (**D**) 22Rv1 (round symbols) and PC3 (square symbols) cells were exposed for 96 hours to LNP-siARNTD (magenta) or LNP-siLuc (yellow). Cell viability was determined by MTS assay. Data represent mean ± SD (n=6) of one representative experiment.

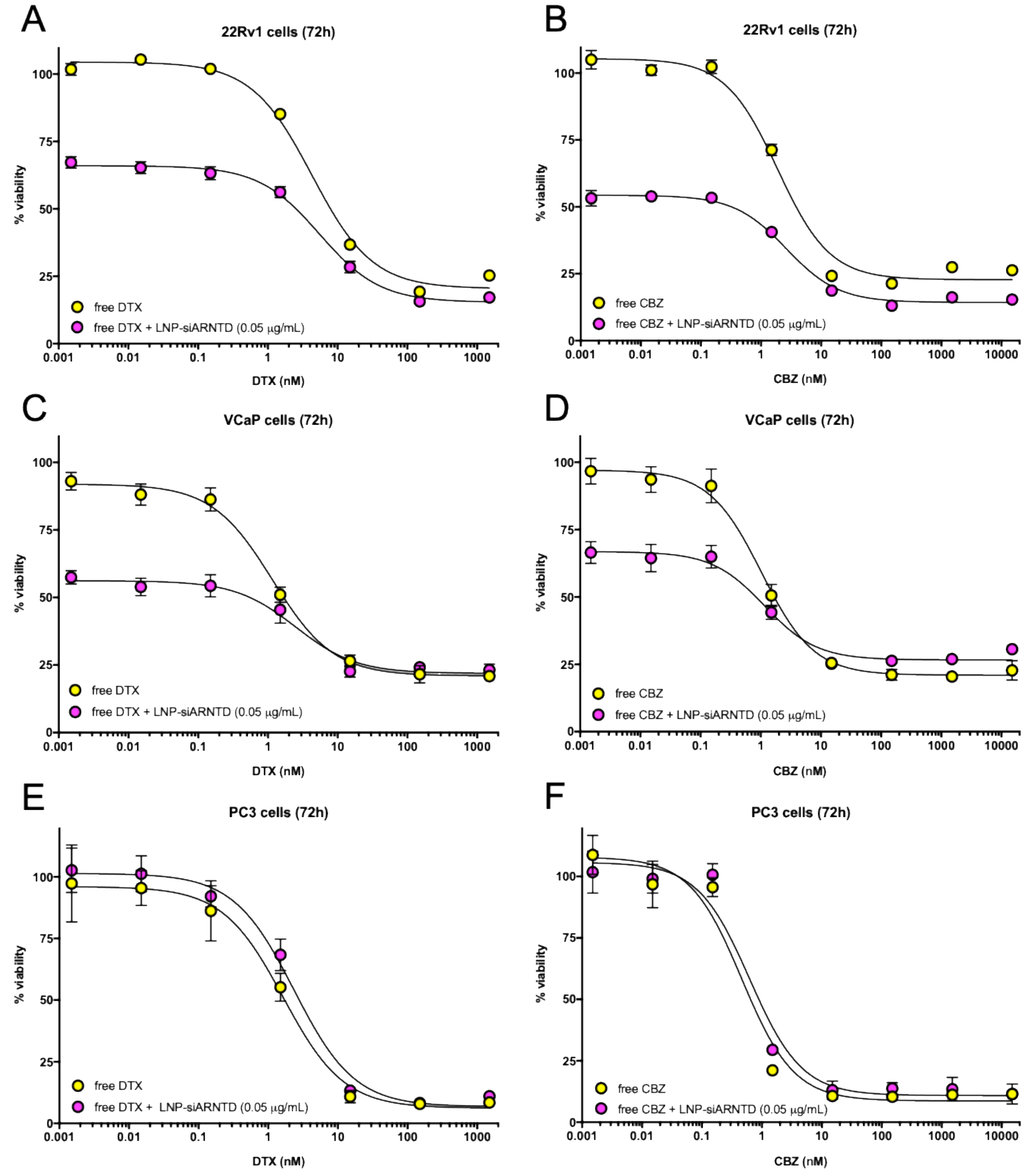


**Supplementary Figure 2. Combining taxane chemotherapeutics and LNPs containing siRNA improves cell viability inhibition**. **(A, B)** 22Rv1, **(C, D)** VCaP and **(E, F)** PC3 cells were exposed for 72h to free docetaxel (DTX) or cabazitaxel (CBZ) in the presence or absence of LNPs containing androgen receptor N-terminal domain siRNA (siARNTD) at a dose of 0.05 µg/mL. Cell viability was determined by PrestoBlue® assay. Data represent mean ± SD (n=6) of one representative experiment.

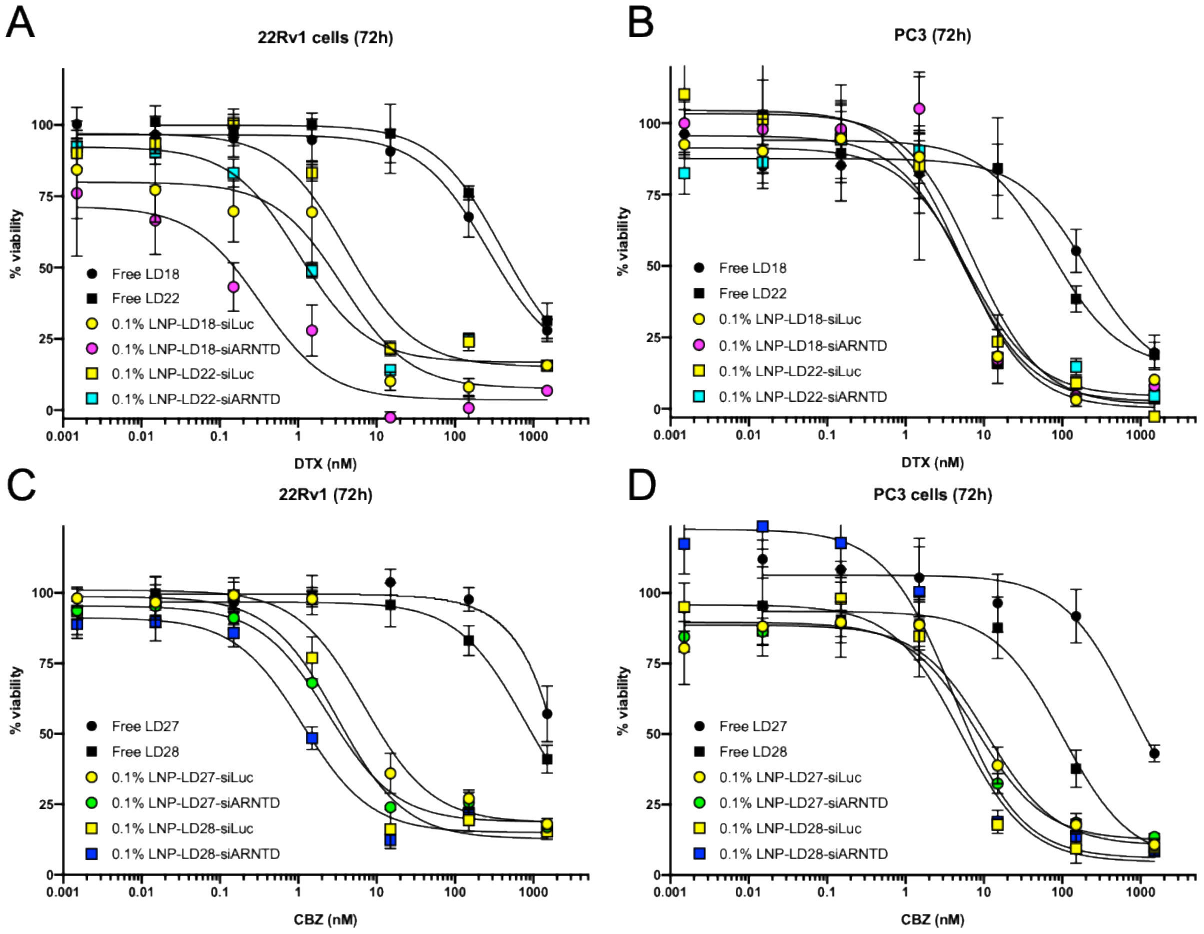


**Supplementary Figure 3. Lipid nanoparticles containing taxane prodrugs and androgen receptor siRNA inhibit cell viability.** (**A, C**) 22Rv1 and (**B, D**) PC3 cells were exposed for 72 hours to LNPs containing 0.1 mol% docetaxel (DTX; LD18, LD22) or cabazitaxel (CBZ; LD27, LD28) prodrug and siRNA against luciferase (siLuc) or androgen receptor N-terminal domain (siARNTD). Cell viability was determined by MTS assay. Data are presented as mean ± SD of one representative experiment (n=6).

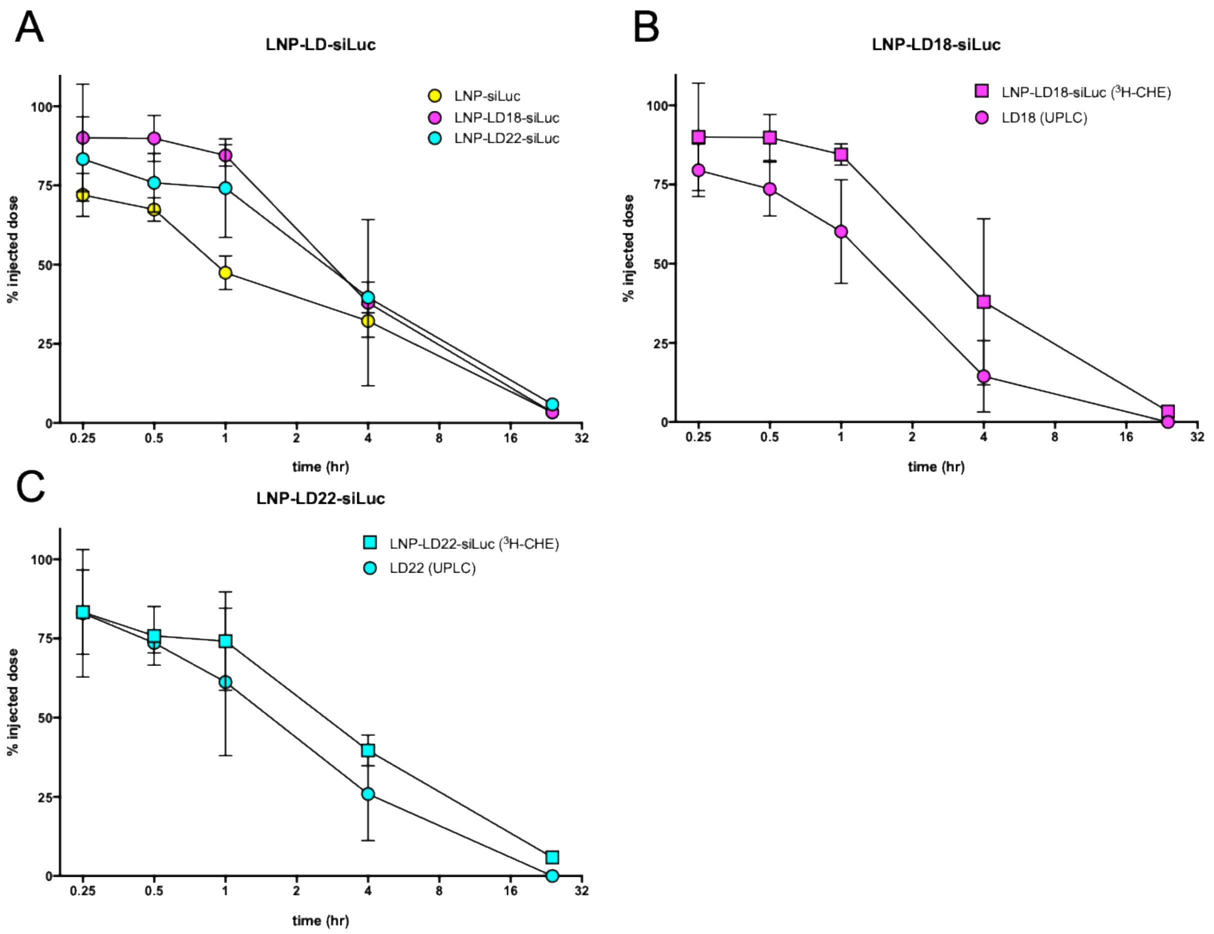


D

| Treatment | $T_{1/2}$ (h) | $V_D$ ((ID)/(%ID)) | CL ((ID)/(%ID)/h) | $AUC^t_0$ (%ID*h) | $AUC^\infty_0$ (%ID*h) |
|---|---|---|---|---|---|
| LNP-siLuc ($^3$H-CHE) | 3.04 | 1.39 | 0.32 | 314.92 | 316.26 |
| LNP-LD18-siLuc ($^3$H-CHE) | 3.06 | 1.00 | 0.23 | 438.04 | 439.96 |
| LNP-LD22-siLuc ($^3$H-CHE) | 3.72 | 1.16 | 0.22 | 457.74 | 463.03 |
| LD18 (UPLC) | 1.59 | 1.11 | 0.48 | 171.43 | 207.89 |
| LD22 (UPLC) | 2.18 | 1.14 | 0.36 | 197.84 | 275.08 |

**Supplementary Figure 4. Pharmacokinetic parameters of radiolabeled LNP formulations following systemic administration in mice.** CD1 mice were intravenously injected with radiolabeled-LNPs containing 10 mol% docetaxel prodrugs at a dose of 2.5 mg/kg siRNA, corresponding to ± 17 mg/kg prodrug. Blood samples were collected via heart puncture at various timepoints and processed for analysis by liquid scintillation counting and UPLC. **(A)** Circulation times of control formulation (LNP-siLuc) and formulations containing 10 mol% LD18 (LNP-LD18-siLuc) or LD22 (LNP-LD22-siLuc). **(B)** Circulation times of LNP-LD18-siRNA and **(C)** LNP-LD22-siRNA as determined by liquid scintillation counting (LNP containing $_3$H-CHE) and UPLC after extraction (LD18 and LD22 prodrugs). Data are presented as mean ± SD of one experiment (n=3-4 animals per timepoint). Pharmacokinetic parameters of LNP formulations determined by PKSolver. $T_{1/2}$, elimination half-life; $V_D$, volume of distribution; CL, clearance; AUC, area under the curve.
